# Intrinsic forebrain arousal dynamics governs sensory stimulus encoding

**DOI:** 10.1101/2023.10.04.560900

**Authors:** Yifan Yang, David A. Leopold, Jeff H. Duyn, Grayson O. Sipe, Xiao Liu

**Affiliations:** Department of Biomedical Engineering, The Pennsylvania State University, University Park, PA, 16802, USA; Neurophysiology Imaging Facility, National Institute of Mental Health, National Institute of Neurological. Disorders and Stroke, and National Eye Institute, National Institutes of Health, Bethesda, MD, 20892, USA; Section on Cognitive Neurophysiology and Imaging, Laboratory of Neuropsychology, National Institute of Mental Health, National Institutes of Health, Bethesda, MD, 20892, USA; Advanced MRI Section, Laboratory of Functional and Molecular Imaging, National Institute of Neurological Disorders and Stroke, National Institutes of Health, Bethesda, MD 20892, USA; Department of Biology, The Pennsylvania State University, University Park, PA, 16802, USA; Institute for Computational and Data Sciences, The Pennsylvania State University, University Park, PA, 16802, USA

**Author notes:** Corresponding Author: Xiao Liu, PhD, 431 Chemical and Biomedical Engineering Building, The Pennsylvania State University, University Park, PA 16802-4400.

## Abstract

The neural encoding of sensory stimuli is subject to the brain’s internal circuit dynamics. Recent work has demonstrated that the resting brain exhibits widespread, coordinated activity that plays out over multisecond timescales in the form of quasi-periodic spiking cascades. Here we demonstrate that these intrinsic dynamics persist during the presentation of visual stimuli and markedly influence the efficacy of feature encoding in the visual cortex. During periods of passive viewing, the sensory encoding of visual stimuli was determined by quasi-periodic cascade cycle evolving over several seconds. During this cycle, high efficiency encoding occurred during peak arousal states, alternating in time with hippocampal ripples, which were most frequent in low arousal states. However, during bouts of active locomotion, these arousal dynamics were abolished: the brain remained in a state in which visual coding efficiency remained high and ripples were absent. We hypothesize that the brain’s observed dynamics during awake, passive viewing reflect an adaptive cycle of alternating exteroceptive sensory sampling and internal mnemonic function.

## Introduction

A core function of the brain is its ability to encode and process external stimuli in both primary and higher-level sensory areas. However, much evidence suggests that the magnitude and reliability of sensory responses in the cerebral cortex and elsewhere depend on its state of arousal (1–4). This dependency is most dramatically evident under different states of consciousness, such as waking, sleep, and anesthesia, when the same sensory stimulus can elicit very different neural responses (2, 3). Even during wakefulness, sensory responses can be highly variable and dependent on momentary arousal level and associated ongoing brain dynamics.

One particularly salient example of arousal state fluctuations affecting sensory responses during wakefulness comes from electrophysiological studies in mice. Cortical responses to visual and somatosensory stimuli are greatly enhanced when mice actively engage in behaviors such as whisking or locomotion, compared to when they are quiescently awake (5–9). Such sensory enhancements generally occur during aroused brain states characterized by desynchronized cortical activity (10–14). In addition, startling an animal with an air puff transiently desynchronizes cortical activity and increases visual responses in the absence of locomotion (15). Similar sensory response effects have been elicited by manipulating the noradrenergic and cholinergic systems, which are thought to underly changes in arousal (16–20). Thus, the arousal state impacts how the brain processes external stimuli during wakefulness. Interestingly, in absence of overt active behaviors and external arousal modulators, a large variability of sensory responses was observed across trials of seconds long(21, 22). This variability has been linked to pre-stimulus ongoing brain activity (21, 22) but might also relate to spontaneous arousal changes (23–25). Nevertheless, the nature and source of the spontaneous seconds-scale fluctuation in ongoing brain activity and arousal remains unclear.

In the absence of external stimuli, the brain exhibits widespread and pronounced spontaneous activity events that transpire over multiple seconds. These coordinated bouts of neural activaty have been observed in both fMRI and electrocorticography (26, 27). Detailed analyses have shown that these neural events usually take the form of relatively slow, spatiotemporal waves that propagate across the cortical hierarchy (28–30) over periods spanning multiple seconds. Similar events have also been observed in large-scale single-unit recordings in the mouse, where temporally coordinated spiking events are evident in cortical and subcortical structures. In fact, up to 70% of neurons from various cortical and subcortical areas are entrained into highly-structured temporal sequences, or cascades, that recur quasi-periodically(27). With each occurrence, individual neurons show spike rate modulation at a consistent phase of the cascade. These observations of large-scale brain activity using different methods appear tightly linked to moment-to-moment changes in central and autonomic arousal levels. For instance, global fMRI peaks and waves are accompanied by deactivation of subcortical arousal-regulating centers and also by changes in delta-band (<4Hz) brain activity (26, 30), whereas spiking cascade is phase coupled to arousal changes measured by pupil size and delta-band (<4Hz) brain activity (27).

While these highly structured, brain-wide events are evident during wakeful rest, it is not known whether they also occur during passive sensory stimulus processing and/or active behavior. If they do, such events might affect the efficacy of stimulus encoding or relate to the known modulation of encoding by locomotion. Importantly, their influence on sensory encoding may intertwine with their established dynamic connection to memory-related neural events(27). Resolving these possibilities will improve our understanding of how the brain balances the distinct tasks of external stimulus processing and internally regulated functions like memory, which have competing demands on neural resources.

To study this phenomenon, we examined spiking activity collected from neuronal populations in 44 cortical and subcortical areas in mice engaged in quiescent rest, visual stimulation, and active locomotion. We found that during periods of visual stimulation in the absence of locomotion, spiking cascades persisted and remained similar to those observed during quiescent rest. Visual responses were systematically modulated across each cascade cycle. The accuracy of visual coding was greatly enhanced in a ∼2 seconds long phase associated with high arousal whereas suppressed in the remaining low arousal phase. Hippocampal sharp-wave-ripple complexes (SPW-Rs), which are critical for memory function (31–33), were oppositely modulated across the high- and low-aroual phases over the cascade cycle. Strikingly, active locomotion abolished cascade sequences and maintained neural activity in the high-arousal phase. Together, these results suggest that highly structured cascades of spiking are entrained by a quasi-periodic, arousal-related cycle that affects activity across the forebrain. We hypothesize that this cycle spurs the alternation two distinct operational modes of the brain, serving sensory and memory functions, respectively.

## Results

To investigate the relationship of intrinsic spiking cascades with visual coding, arousal, and hippocampo-cortical dynamics, we used the Visual Coding – Neuropixels dataset from the Allen Institute (34, 35). We analyzed spiking activity of ∼22,000 neurons recorded from 32 mice (683 ±140 neurons per mouse, mean ±SD, see **Table S1** and Methods for details), spread across 44 brain regions, mostly in the visual cortex, hippocampus, and thalamus (**Fig. 1A**). We focused on 7 experimental sessions, including 3 involving natural scene visual stimuli, 3 involving drifting-gratings, and one spontaneous session without stimulation (**Fig. S1A**).

**Figure 1.**
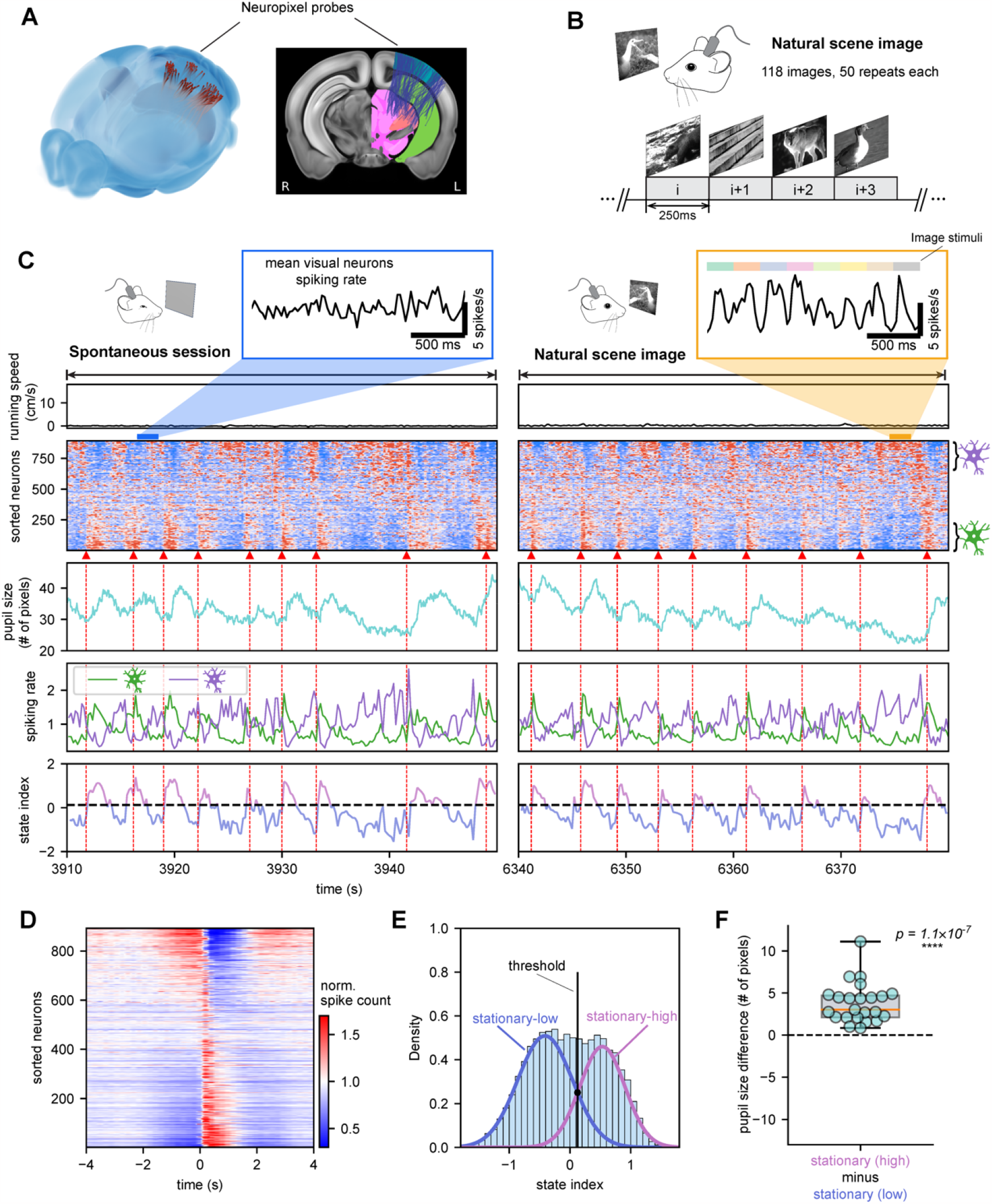
Spiking cascades of arousal relevance persist during continuous visual stimulation. (A) Locations of 183 neuropixel probes from 32 mice in the Allen Mouse Brain Common Coordinate Framework (version 3). Left: the 3-D representation of probe insertion in mouse brain. Right: the 2-D projection of the probes onto a middle brain slice highlights the major recording sites: visual cortex (blue), hippocampus (green) and thalamus (pink). (B) Illustration of natural scene image session where grayscale images were continuously presented to mice for 250ms each in random order through a monitor. The 118 images were presented for 50 times each, across 3 different sessions in every mouse. (C) Example data from a representative mouse under two different conditions. Left column: A 40-s stationary period of spontaneous session without stimulation. In the absence of running (first row), the neuronal spikes were organized in the form of spiking cascade (second row: neurons were sorted by the principal delay profile derived by a data-driven method) with sequential activation from the negative-delay neurons (fourth row: purple colored) to the positive-delay neurons (fourth row: green colored). The spiking cascade is phase-coupled to changes in pupil diameter (third row). A state index was then defined based on the relative activation level of positive-delay neuron and negative-delay neurons with pink / blue colors denoting the high / low arousal states respectively (fifth row). Right column: A 40-s stationary period of the nature image stimulation session that showed similar spiking cascades during visual stimulation. The pop-up panels at the top show the mean spiking rate of all visual neurons in example 2-sec time segments, which was phase-locked to stimuli in the image stimulation session (orange) and absence of systematic pattern in the spontaneous session (blue). See **Fig. S5** for more detail. (D) The average pattern of the spiking cascade in the representative mouse. (E) Distribution of state index in stationary periods (light-blue-colored) from a representative mouse. The stationary distribution is fitted with a two-classes gaussian mixture model, yielding the fitted curves for the stationary-high state (pink-colored) and stationary-low state (blue-colored), as well as the optimal boundary between the two (black dash line). (F) The boxplot of pupil diameter difference between the stationary-high and stationary-low states for mice with pupil data. Each dot represents a mouse and the one-sample t-test (two-sided, N=23) is used for significance test.

### Spiking cascades pervade the cortex during visual stimulation

We first ordered simultaneously recorded neurons based on their principal delay profile, as per previously established convention (27). We compared periods with and without visual stimulation, both in absence of locomotion (“stationary behavior”) (**Table S2**). Against expectation, the cascades of spiking activity pervaded brain activity in a manner that was both qualitatively and quantiatively similar between the two conditions (**Fig. 1C**, see also **Figs. S2, S3**). Within each cascade, individual neurons had a fixed temporal relationship to one another, with the average over all cascades in a representative session shown in **Fig. 1D**. The similarity of cascade dynamics with and without visual stimulation is reflected in highly similar (*r* = 0.836 ± 0.08, mean ±SD; *p* = 8.5×10^−84^**; Fig. S1D**) principle delay profiles across the two conditions.

We defined a state index summarizing the relative activation level of the negative- and positive-delay neurons, the neuronal subpopulations active during distinct phases of each cascade cycle (**Fig. 1C** and **1D**; **Fig. S4**). The state index served as a proxy for the internal state of the brain entrained to each cascade. It showed a bimodal distribution during the stimulated stationary periods, suggesting the existence of two distinct brain states (**Fig. 1E**). Based upon the reliable variation of the state index with pupil size changes (**Fig. 1F**), we refer to the two states low-arousal and high-arousal, which correspond well to distinct phase of the cascades (bottom row in **Fig. 1C**). During high arousal state, the pupil was dilated relatively and positive-delay neurons showed their highest activity, whereas during low arousal state the pupil was constricted and negative-delay neurons showed their highest activity.

These findings demonstrate that the spiking cascade dynamics observed previously during quiescent, unstimulated rest persist in approximately the same form during periods of continuous and intense visual stimulation when the animal is stationary. These apparently self-generated dynamics, which are not time-locked to much more rapid (i.e., 250 ms) external stimulus presentations, cause the brain to engage in quasi-periodic fluctuations with intervals of 5-10 seconds. Within each cycle, the brain alternates between two distinct states that vary in their arousal levels.

### Stimulus encoding is systematically modulated during each cascade cycle

We next investigated how the cascade dynamics may affect the encoding of visual stimuli among recorded neuronal population. To study this, we employed machine learning models to measure the precision of visual information encoding and its dependency on neural population dynamics. We trained support vector machine (SVM) decoders with a 5-fold cross validation to predict the image identity based on the neuronal spiking activity it evoked and recorded the correctness of the prediction for each of the 118 stimuli × 50 repeated presentations (**Fig. 2A**, also see **Methods** for details). The prediction accuracy of the decoders serves as a measure of encoding performance across different conditions and reflects the efficacy of visual sensory processing.

**Figure 2.**
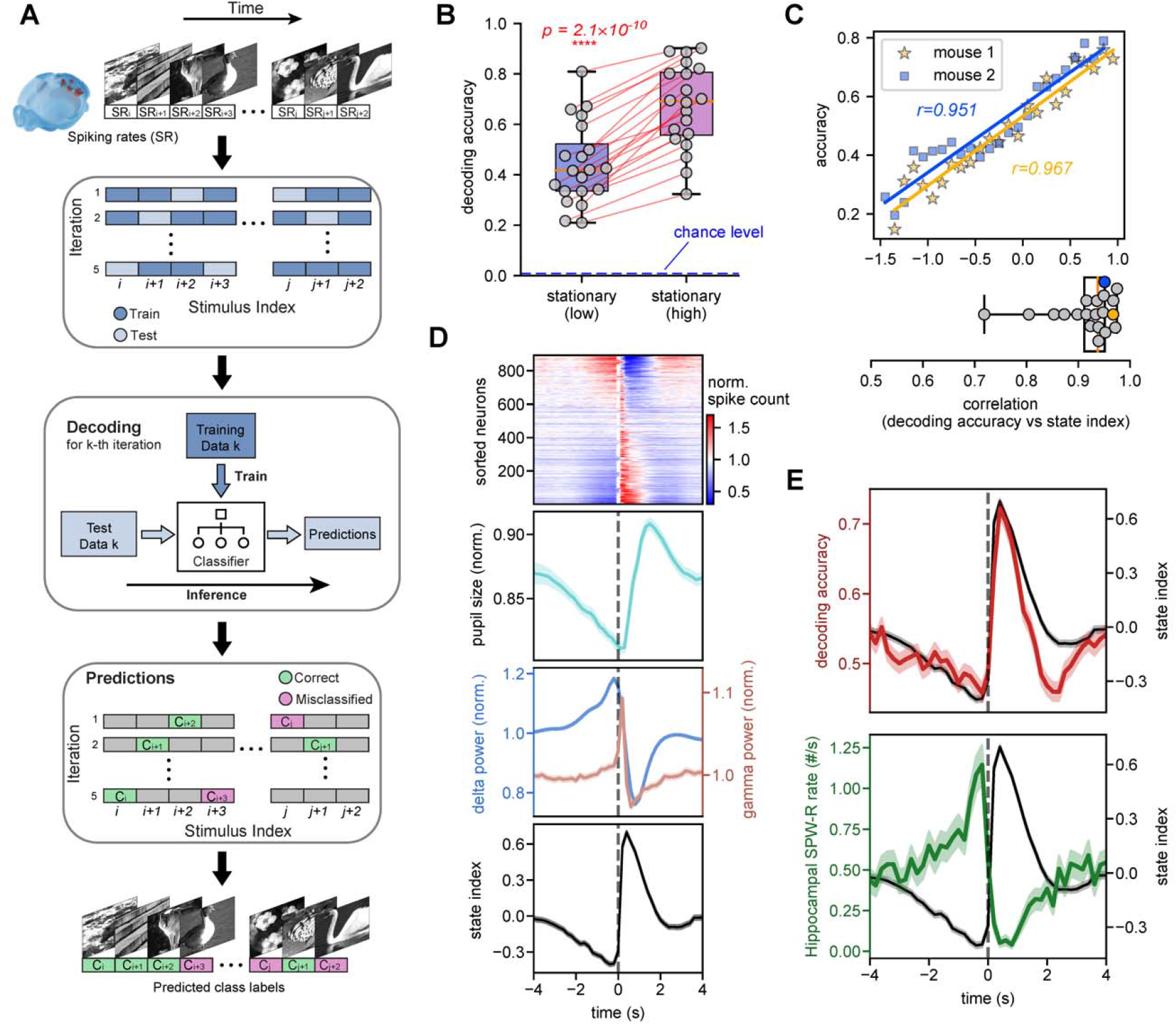
Population encoding of visual stimuli tightly follows the cascade dynamics. (A) Pipeline for training a machine learning model to predict visual stimuli based on population spike data. The 118X50 stimulation trials (250 ms, 50 repetitions for each of 118 image stimuli) were shuffled and evenly split into 5 groups with each covering 10 repetitions of all the 118 stimuli. For each iteration, an SVM-based decoder was trained with data of 4 groups and then used to make inference on the remaining group. Every trial was evaluated and labeled either correctly classified or misclassified after 5 iterations. (B) Box plot showing the decoding accuracy summarized for different conditions. Each dot represents a mouse and the two-sided pairwise t-test was used for significance test (N=20 mice). The blue dashed line marked the chance level of 0.0085. (C) The linear relationship between state index and decoding accuracy is shown for two example mice (Top) and also summarized in a box plot for all mice (Bottom), where the yellow and blue dots represent the example mice. The linear relationship was derived from the data during the stationary periods of continuous visual stimulation. (D) Various behavior and neuronal signals averaged over the 8-s cascade cycle, from top to bottom showing the averaged cascade pattern, pupil size, LFP band powers, and the state index. Note the averaged cascade pattern is from the representative mouse. (E) The opposite modulations of the decoding accuracy (top) and hippocampal sharp wave ripples (SPW-Rs) rate (bottom) across the cascade cycle during stationary periods with natural image stimulation. All of the time series data is shown as mean ± SEM from all cascade events (N=1401) across all 32 mice.

The encoding of visual stimuli was strongly affected by the internal state defined by population spiking dynamics. On average, the decoding accuracy showed a striking difference (23.0 ± 8.5%, *p* = 2.1×10^−10^, *N* = 20 mice, paired t-test) between the two internal states defined through population spiking activity (**Fig. 2B**). Given that the two stationary states were defined based on the state index, we directly examined the relationship between the state index and decoding accuracy. This analysis revealed a strong and robust linear association (*r* = 0.975, *p* = 3×10^−21^; all mice data; **Fig. S6**) between the two, which was highly reproducible across individual mice (**Fig. 2C**).

During stationary periods of visual stimulation, these different modes of stimulus encoding were attached to different phases of the cascade cycle. The encoding accuracy was temporally entrained, along with multiple arousal measures, to the occurrence of each cascade event (**Fig. 2D** and **2E**). Arousal measures such as pupil size, delta (<4Hz) and gamma (55-65Hz) local field potential (LFP) power, and the previously computed state index, were systematically modulated with each cascade cycle (**Fig. 2D**). In particular, the decoding accuracy and state index were modulated in a highly similar way, with their fast descending phase (0.3-2 seconds with respect to the cascade center) coinciding actually with the pupil dilation. Importantly, simultaneous measurements from the hippocampus indicated that sharp wave ripples (SPW-Rs) were modulated in an antagonistic way to both: the ripple rate was the highest at the troughs of state index and decoding accuracy whereas it decreased to almost zero at state index peaks(**Fig. 2E**).

We next investigated the effect on visual encoding in multiple different visual regions of the mouse brain, while using the hippocampus as a control region. Performance of the decoder was able to draw upon all visual areas, with the lateral geniculate nucleus (LGN) and primary visual cortex (VISp) showing the strongest contribution, such that their exclusion led to the strongest reduction in decoding accuracy (**Fig. 3B**). Analyzing of the weight contribution in the SVM models (**Fig. S7A**) and the accuracy of decoders trained with single-region data (**Fig. S7B**) yielded similar results.

**Figure 3.**
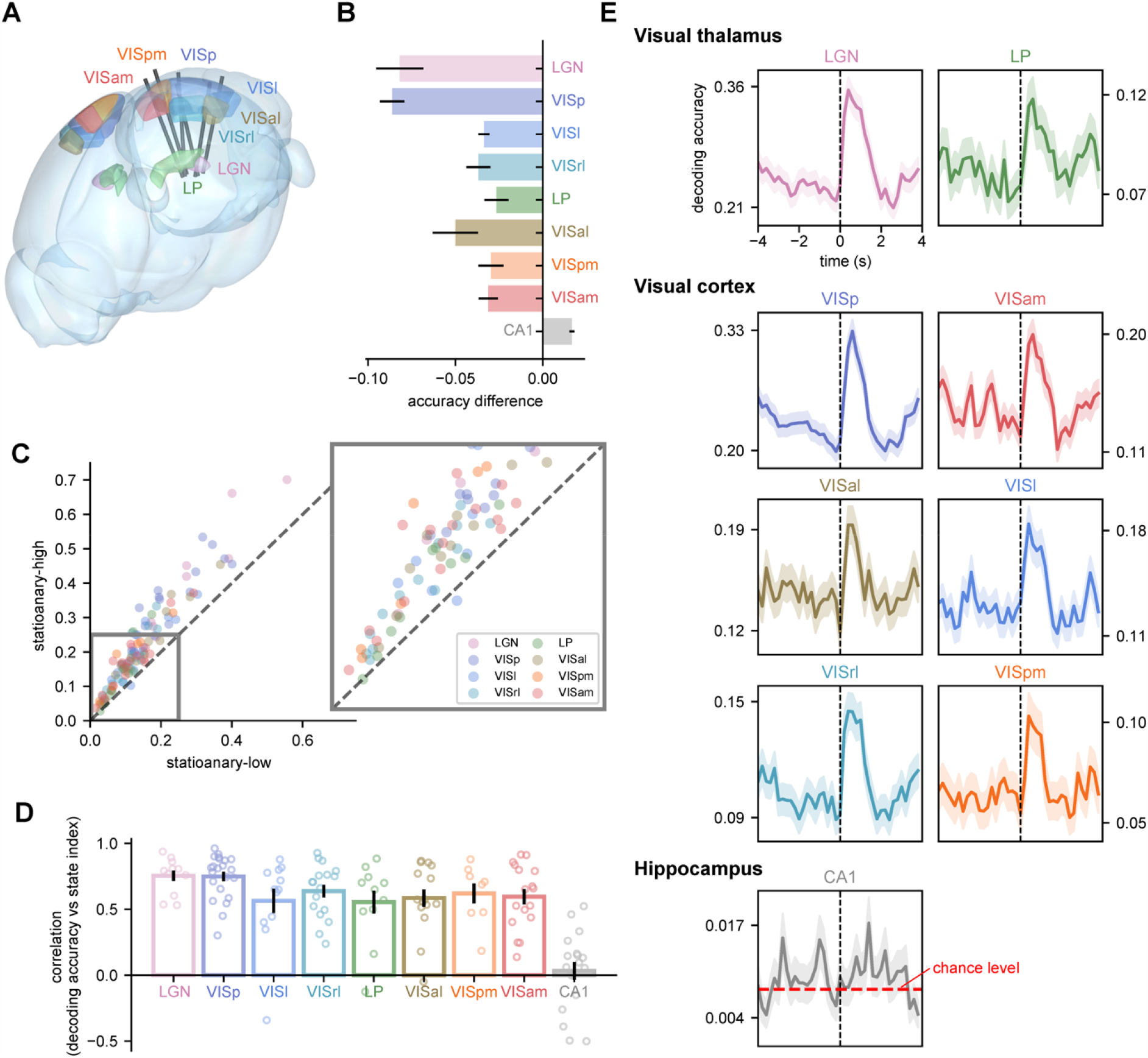
Cascade phase dependent visual coding is evident in all visual areas. (A) Schematic layout of Neuropixel probes relative to their target visual regions, including primary visual cortex (VISp) and five high-order visual cortical areas, i.e., latero-medial area (VISl), anterol-ateral area (VISal), rostro-lateral area (VISrl), postero-medial area (VISpm) and antero-medial area (VISam) (image credit: the Allen Institute). The probes with insertion depth up to 3.5mm into the brain also record the spiking activity within two visual thalamic nuclei, i.e., the lateral posterior nucleus (LP) and the lateral geniculate nucleus (LGN), as well as other regions traversed by the probes, e.g., the hippocampus. (B) A bar plot showing the change of decoding accuracy resulting from excluding a specific region. (C) A comparison of the accuracy of region-specific decoders, which were trained with data of a specific brain region, between the stationary-high and stationary-low states. Each colored dot represents a specific visual region from a single mouse. (D) A box plot showing the linear relationship between the state index and the accuracy of region-specific decoders, similar to **Fig. 2C**. Each dot represents a mouse with the corresponding region recorded. (E) The region-specific decoding accuracy over the 8-s spiking cascade cycle from all the mice with the corresponding region recorded (N=1401). The red dashed line marked chance level equal to 0.0085.

Importantly, the accuracy of each single-region decoder was consistently higher during the stationary-high state than the stationary-low state (**Fig. 3C**) and actually correlated with the state index (**Fig. 3D**), indicating that neurons in each of the areas were affected by the cascade fluctuations in qualitatively similar ways. This is also evident in the dynamics of each area’s encoding accuracy time course over the cascade cycle (**Fig. 3E**). These results were consistent across different choice of decoder models, i.e., logistic regression (**Fig. S8**) and multilayer perceptron (**Fig. S9**). In each case, shuffling the stimulus labels completely abolished decoder performance (**Fig. S10**).

### Locomotion abolishes cascade dynamics during visual presentation

Given previous findings that locomotion affects visual information processing (5, 9, 15, 19), we investigated whether the cascade dynamics and visual coding accuracy would be affected by the mouse’s running behavior. We found that, during running episodes, spiking cascades nearly vanished and the cyclical nature of arousal seen in absence of locomotion ceded to a continuous condition of high-arousal, with negative-delay neurons continually suppressed whereas the positive delay neurons showing sustained activation (**Fig. 4A**). Hippocampal ripples were nearly abolished (**Fig. 4B**). In contrast to the stationary periods of visual stimulation, the computed state index during active running and visual stimulation had a single peak in its distribution (light yellow, **Fig. 4C**), corresponding to the stationary-high arousal state. Accordingly, visual coding accuracy during running was much higher than the stationary periods overall (**Fig. 4E**, *p* = 1.9×10^−5^, paired t-test, *N =*23 mice) but similar to the stationary-high state (*p =* 0.13, paired t-test, *N =*23 mice). Neverthless, the running periods were associated with much larger pupil size than the stationary-high periods (**Fig. 4D**, *p* = 2.4×10^−13^, paired t-test, *N =*23 mice).

**Figure 4.**
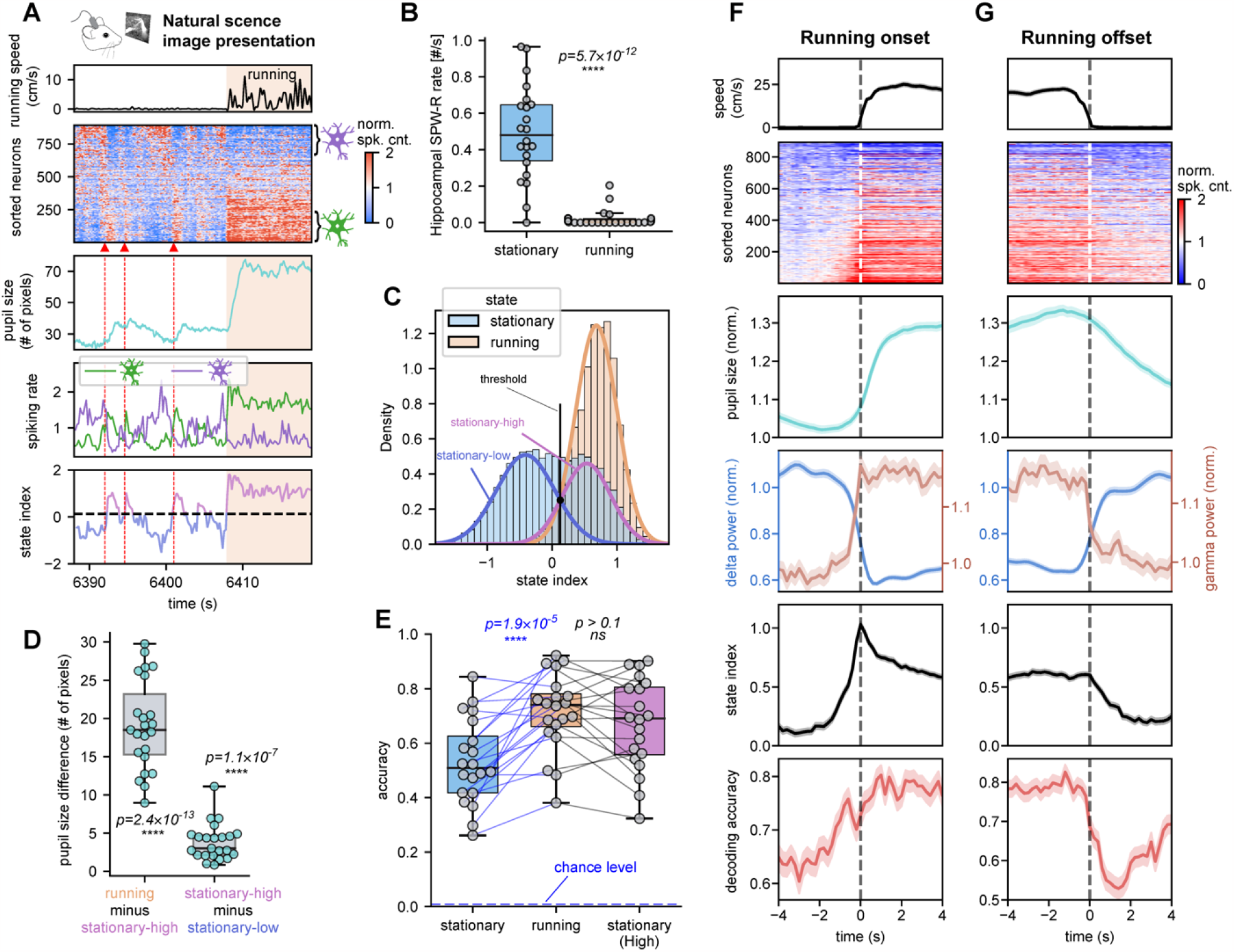
Running sustains visual coding efficiency and abolishes hippocampal sharp-wave ripples. (A) A 30-s period of the natural image session at the transition to a running episode. The transition is accompanied by the cessation of spiking cascades, a large increase in pupil diameter, sustained activation / deactivation of positive-delay / negative-delay neurons, and a high state index. (B) The occurance rate of the hippocampal SPW-R summarized for stationary (light-blue) and running (orange) respectively, which are significantly different from each other (two-sample t-test, two-sided). (C) The state index distribution of the representative mouse during running periods (orange) as compared with the stationary periods (light blue). (D) The boxplot of pupil diameter differences for 23 mice with pupil data. The averaged pupil diameter differences were separately summarized between the running and the stationary-high state periods, and between the stationary-high and stationary-low states. Each dot represents a mouse and the one-sample t-test (two-sided, N=23 mice) is used for significance test. (E) The decoding accuracy summarized for the running peirods and the two stationary states. Each dot represents a mouse and the pairwise t-test was used for significance test (two-sided, N=20). (F) Changes of behavior and neuronal signals and visual coding accuracy at the running onset. The data were averaged over all onset events (N=465) during natural scene image session across all 32 mice, from top to bottom showing the running speed, averaged cascade pattern, pupil size, delta/gamma power, state index, and neural decoding accuracy. (G) The change of signals at the running offset (N=401), same as in (F).

Given this clamping to a high-arousal state during periods of locomotion, the onset and offset of running provided an additional perspective on the temporal dynamics of the associated brain state changes. We found that decoding accuracy and state index preceded running onset by ∼ 1.5 sec and took ∼1.5 sec to slowly decrease back to the baseline after the cessation of running (**Fig. 4F** and **4G**). The pupil dilation/constriction showed similar, albeit sluggish, changes at both running onsets and offsets.

### Cascades affect visual coding by modulating the magnitude of sensory responses

To understand the basis of changes in population encoding of visual information, we examined how single-neuron responses are modulated by the cascade dynamics. For this purpose, we focused on the drifting-grating stimulation sessions, which include a 1-sec baseline period between stimulations and thus enable the estimation of evoked spiking response amplitudes (**Fig. 5A**). Confirming the findings above, we found that the orientation and temporal frequency of drifting-grating stimuli can be more accurately decoded from the neuronal data of the stationary-high state than that of the stationary-low state (**Fig. S11**). Inspecting single-neuron responses, we identified two groups of neurons showing significant but opposite responses to the drifting-grating stimuli (enhanced and suppressed neurons in **Fig. 5B**). Neurons may respond to more than one grating direction, but the type of response (enhanced or suppressed) was largely consistent (**Fig. S12A**).

**Figure 5.**
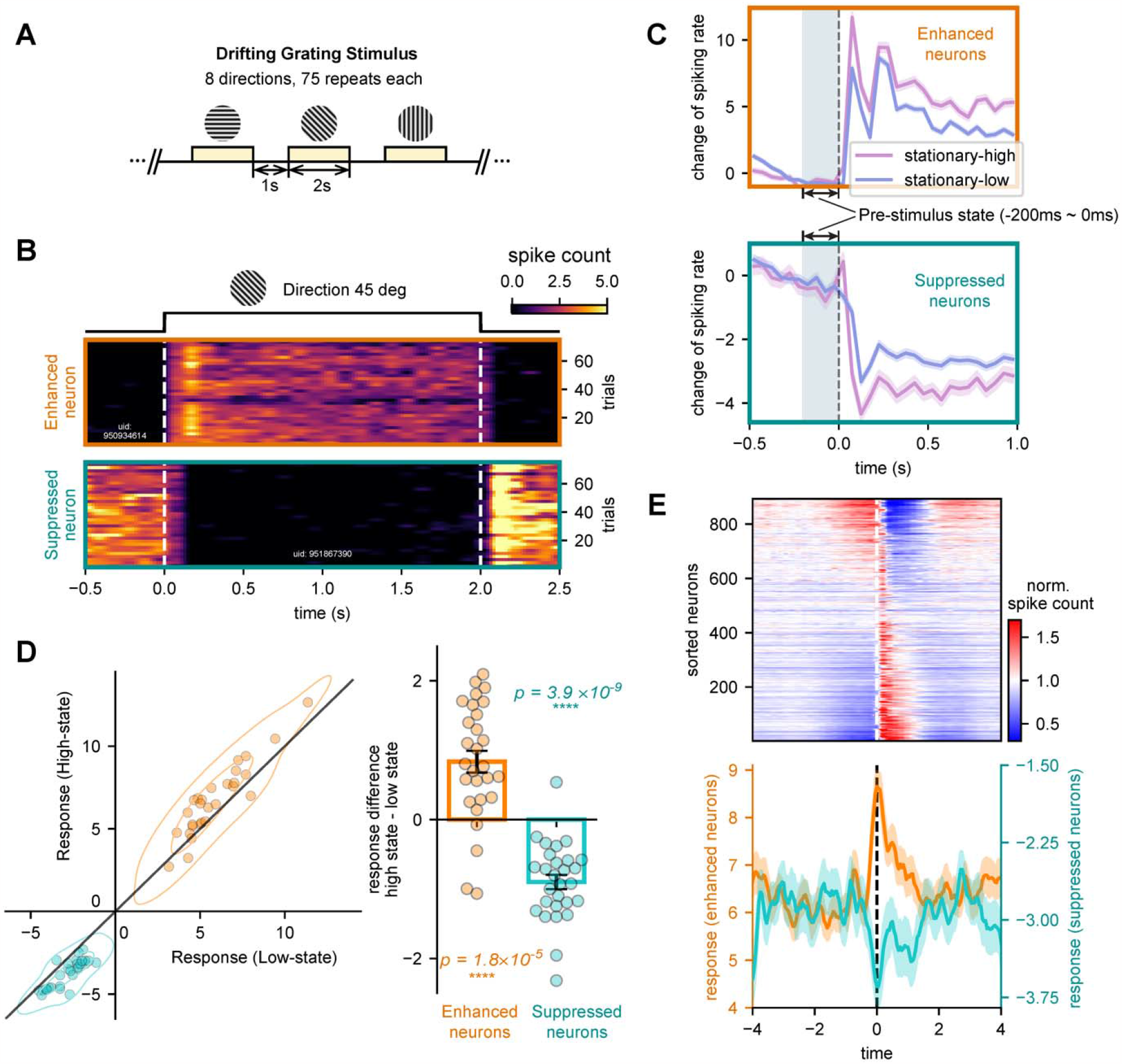
Spiking cascade modulates single-neuron responses to change signal-to-noise ratio. (A) Illustration of stimulus presentation in the drifting-grating session. Oriented drifting gratings were repeatedly presented to mice for two seconds following a one-second blank period. (B) Examples of two types of neurons (top: drifting grating enhanced neuron; bottom: drifting grating suppressed neuron) that showed opposite responses to a drifting grating stimulus. (C) The mean responses of enhanced neurons (top) and suppressed neurons (bottom) in a representative mouse differ with respect to the pre-stimulus state (−200 to 0ms before stimulus onset) defined by the state index. The results are shown as mean ± SEM. (D) Response amplitude compared between the trials with different pre-stimulus states for both enhanced (orange) and suppressed (teal) neurons. Single neuron response is quantified as the mean spiking rate difference between stimulus onset (0 to 400ms) and baseline (−800 to 0ms). Each dot represents a mouse, the one-sample t-test is used for significance test (two-sided, N=28). (E) The mean responses of the enhanced (orange) and suppressed (teal) neurons are dependent on the phase of their pre-stimulus period in the cascade cycle (bottom). The 160 time bins between -4 sec to 4 sec have different numbers of trials ranging between 82 to 168. The averaged cascade pattern (top) is from the representative mice.

We next investigated how the brain state affects single neuron responses. Although the drifting-grating responding neurons showed no systematic overlap with the negative- and positive-delay neurons of the cascade, we excluded the drifting-grating responding neurons classified also as either negative- or positive-delay neurons for subsequent analyses to avoid any potential confounding (**Fig. S12B**). We found that the pre-stimulus state (−200ms−0ms) significantly affected the amplitude of evoked responses. The stimuli presented right after the stationary-high state elicited larger reponses of the enhanced neurons but decreased spikes of the suppressed neurons to a larger extent, compared with those presented right after the stationary-low state. This is evident in trial averages (**Fig. 5C**) and across individual mice (**Fig. 5D**). A similar result was also observed between the running and stationary periods (**Fig. S12C and S12D**). Importantly, after sorting the trials according to the location of their pre-stimulus period in the cascade cycle, the response of the drifting grating enhanced and suppressed neurons clearly showed opposite modulations across the cascade cycle in a pattern similar to what we observed with population decoding accuracy (**Fig. 5E**). These changes in single-neuron responses could potentially affect the signal-to-noise (SNR) ratio of visually evoked responses, and thus influence the efficacy of visual coding over the cascade cycle.

## Discussion

Here we show that the mice brain, during stationary periods of passive visual stimulation, showed quasi-periodic state fluctuations indicated by stereotyped population dynamics spanning several seconds. These dynamics manifest as cyclical spiking cascades entraining the activity of a high proportion of neurons in the cortex, hippocampus, and thalamus. These cascades are accompanied by changes in hippocampal ripple activity and arousal indictors, such as pupil size. Interestingly, the cascade strongly influences the brain’s capacity to accurately encode visual information, in a manner opposite to its effect on the hippocampal ripples. Moreover, locomotion abolished both the cascade cycles and hippocampal ripples, leading to prolonged periods of high arousal and efficient visual encoding. The results thus cast new light on the nature of neural variability in behaving animals.

We show that ongoing dynamics responsible for the moment-to-moment variability in neural responses are intricately organized and involve neural populations distributed across the brain. It is generally appreciated that an identical stimulus can lead to different neuronal responses depending on brain sate (1, 5, 11, 15, 36). This variability can play out over multiple time scales, including over a period of several seconds (15, 21, 25). Previous efforts to link this response variability to ongoing pre-stimulus activity (21, 22) have not focused on how the brainwide neural dynamics might contribute to the observed variability in local responses (37, 38). While arousal fluctuations are recognized as likely contributing to variable sensory responses (23, 27), the precise underlying network mechanisms are still poorly understood.

The discovery of cyclical cascades pervading the brain during rest (27) suggests that arousal fluctuations are fundamentally related to the phase of these cascades. Thus, in contrast to previous studies seeking to link sensory responsonse variability to arousal fluctuations measured by pupil diameter changes (23–25), we focused on the role of cyclical brain-wide cascades on population visual encoding. Our discovery of persisting cascades during periods of visual stimulation, and their striking relationship with sensory encoding, indicate a fundamental signature and perhaps mechanism governing brain states. These findings have implications for understanding the neural basis of similar large-scale brain dynamics measured using other methods such as fMRI and ECoG, which have been described as infraslow waves propagating along a cortical hierarchical gradient (28–30, 39). Like the cyclical spiking cascades measured in the present study, these propogating waves are also coupled to arousal fluctuation on the timescale of seconds, and could be the macro-scale representation of cascade dynamics.

The influence of endogenous cascade dynamics over sensory encoding may have important functional implications. For example, alternating between modes of efficient stimulus encoding and hippocampal ripple activity may be relevant for mechanisms of learning and memory. Previous work has proposed that alternation of distinct brain states serves memory consolidation across sleep (40), with cholinergic neuromodulation being a central component. According to this view, elevated cholinergic tone characteristic of wakefulness, prioritizes the encoding of new information into the hippocampus, whereas a reduced tone during sleep favors memory consolidation from the hippocampus to the cortex (40). Increased cholinergic and adrenergic modulatory activity is indeed correlated with spontaneous pupil dilations (25, 41) and enhanced excitability of visual neurons (17, 41–44), which are similar to what we observed at the high-arousal phase of the cascade. The endogenous cascades we describe here may therefore represent the same alternation between sensory encoding and mnemonic function but on shorter timescales and entirely during wakefulness. The intrinsic cascades could thus drive the seconds-scale oscillation between two modes during wakefulness even under continuous sensory stimulation. Such quick and repeated switching between the two functional modes, which mimics the alternating of information forwarding and error back-propagating during the learning of artificial neuronal networks (45), might be generally important for learning and memory.

Another conjecture on the cascade function is related to its sequential nature, in which neurons activated at similar phases are more synchronized. Individual cascades might serve as a reference event to help bind information presented closely in time. The new finding of the correspondence between the cascade sequence and the hippocampal replay sequence supports this hypothesis (46). Additional experiments are needed to test this exciting possibility.

Finally, our results suggest that the locomotion-related visual coding enhancement observed in previous studies (5, 9, 15) is enabled by the accompanying highly aroused brain state, rather than the act of locomotion itself. In examining the initiation and termination of running bouts, we found that the associated brain state dynamics, rather than the actual locomotion, promoted a sustained period of high visual encoding accuracy with virtually devoid of hippocampal ripples. The locomotion-related state change thus appeared to replace the intrinsic cascade dynamics with a continuously high arousal phase that prioritizes information encoding. Our findings may also be consistent with previous work suggesting that relatively low forebrain arousal is a prerequisite for optimal memory function (47, 48). During sustained periods of high arousal, such as locomotion, the brain has diminished internal mnemonic processes and thus deteriorated memory function compared to stationary modes of wakefulness. It is interesting to consider that memory consolidation, rather than being relegated to offline processing during sleep, is an active and prominent feature of the awake brain, but only during some behavioral states. From this perspective, the brain-wide spiking cascades investigated in the present study may be central mediators of this process, periodically switching between endogenous cycles of sensory sampling and mnemonic engagement.

## Acknowledgements

This work was supported by the Brain Initiative award (1RF1MH123247-01), the NIH R01 award (1R01NS113889-01A1), and the Intramural Research Program of the National Institute of Mental Health (ZIA-MH002838).

## Author Contributions

**Y.Y.&X.L**. contributed to the conception, design of the work, and data analysis;

**X.L**. also devoted the efforts to the supervision, project administration and funding acquisition;

**Y.Y**., **D.A.L**., **J.H.D. G.O.S. & X.L**. contributed to data visualization, and writing the paper.

## Competing interests

Authors declare that they have no competing interests.

## Data and materials availability

We used the Neuropixels Visual Coding dataset from the Allen Institute (34, 35). All the multimodal data are available at https://portal.brain-map.org/explore/circuits/visual-coding-neuropixels. The Python code that produced the major results of this paper will be available at https://github.com/psu-mcnl/Neural-Arousal.

## Notes

https://portal.brain-map.org/explore/circuits/visual-coding-neuropixels

### Competing Interest Statement

The authors have declared no competing interest.

