## Supplementary Materials for "Intrinsic forebrain arousal dynamics governs sensory stimulus encoding"

### 1 Supplementary Tables

#### 1.1 Table S1

**TableS1 Mice exclusion details.**

| | # of Mice removed | # of Mice remained | Removal details |
|---|---|---|---|
| Initial removal | 3 | 29 | Mice with insufficient stationary periods (≤10%) were removed.<br>• 758798717: 0.3% stationary periods<br>• 760693773: 1.8% stationary periods<br>• 762602078: 0.0% stationary periods |
| Pupil analysis | 6 | 23 | Mice with no pupil data available were removed.<br>• 715093703<br>• 719161530<br>• 721123822<br>• 732592105<br>• 737581020<br>• 739448407 |
| Natural scene visual stimuli decoding analysis | 9 | 20 | Mice with insufficient number of samples (n≤150) for any of the condition (stationary-high, stationary-low, running) were excluded.<br>• 732592105: 76 samples for stationary-high.<br>• 737581020: 141 samples for stationary-high.<br>• 746083955: 81 samples for stationary-high.<br>• 757216464: 0 samples for both stationary-high and low.<br>• 760345702: 0 samples for both stationary-high and low.<br>• 761418226: 0 samples for both stationary-high and low.<br>• 762120172: 48 samples for stationary-high.<br>• 791319847: 127 samples for stationary-high.<br>• 798911424: 147 samples for stationary-low. |
| drifting-gratings response analysis | 1 | 28 | Mice lacking stationary periods during the drifting-grating sessions were excluded.<br>• 75134857: 0.0% stationary periods during drifting-grating sessions. |

#### 1.2 Table S2

**Table S2 Stationary/running periods summarized for stimulus sessions across all mice.**

| Sessions | Stationary periods (sec) | Stationary percentage (%) | Running periods (sec) | Running percentage (%) |
|---|---|---|---|---|
| Nature image | 542.4 ± 457.2 | 44.3 ± 34.0 | 625.8 ± 409.3 | 55.7 ± 34.0 |
| Drifting-gratings | 863.5 ± 559.0 | 56.1 ± 31.1 | 622.3 ± 461.6 | 43.9 ± 31.1 |
| Spontaneous | 552.6 ± 321.8 | 55.3 ± 27.9 | 412.4 ± 254.8 | 44.7 ± 27.9 |

* Mean ± SD

### 2 Supplementary Figures

#### 2.1 Figure S1

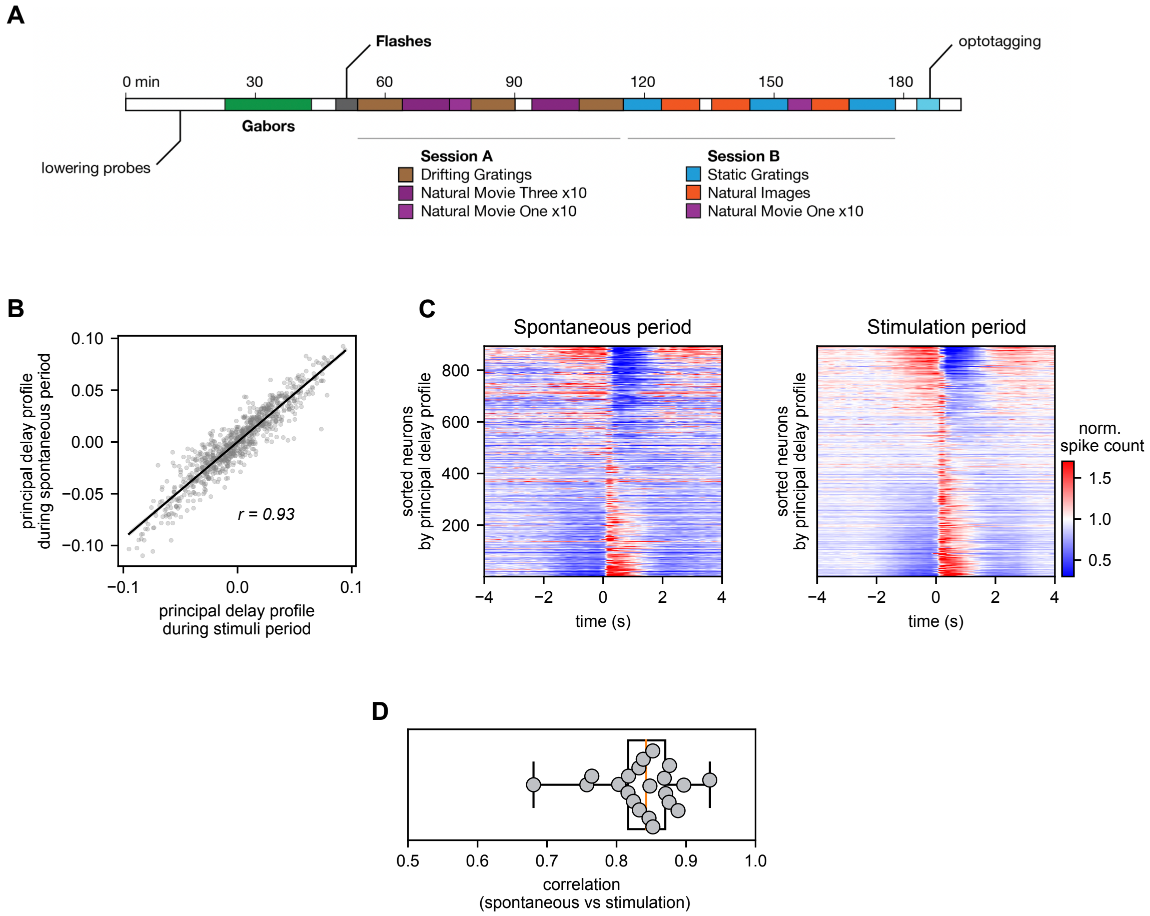

**Figure S1**. Spiking cascades pervade the cortex during both visual stimulation and rest. (A) Illustration of the "Brain Observatory" stimulus set from Allen Visual Coding - Neuropixel dataset. (B) Assessment of the similarity between the principal delay profile obtained from spontaneous neural activity and that from neural activity during visual stimulation in a representative mouse. The degree of similarity is quantified using Pearson's correlation coefficient. (C) The average pattern of the spiking cascade during spontaneous period (Left) and visual stimulation period (Right) from the representative mouse. (D) Box plot illustrating the correlation between principal delay profiles of spontaneous and visual stimulation sessions across individual mice. Each data point represents a distinct mouse.

#### 2.2 Figure S2

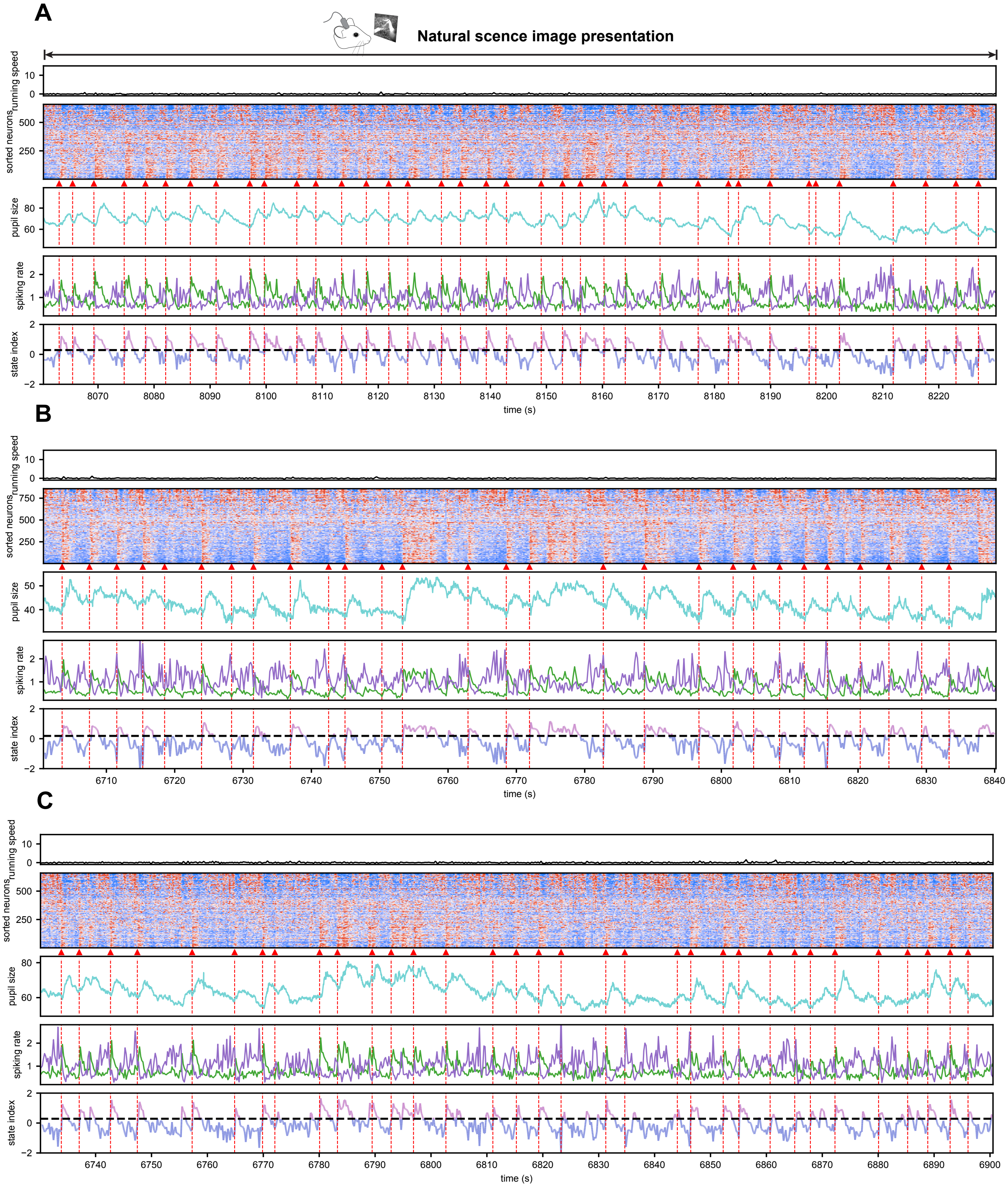

**Figure S2**. (A)–(C) Examples of spiking cascade during continuous natural scene image stimulation in the absence of running.

#### 2.3 Figure S3

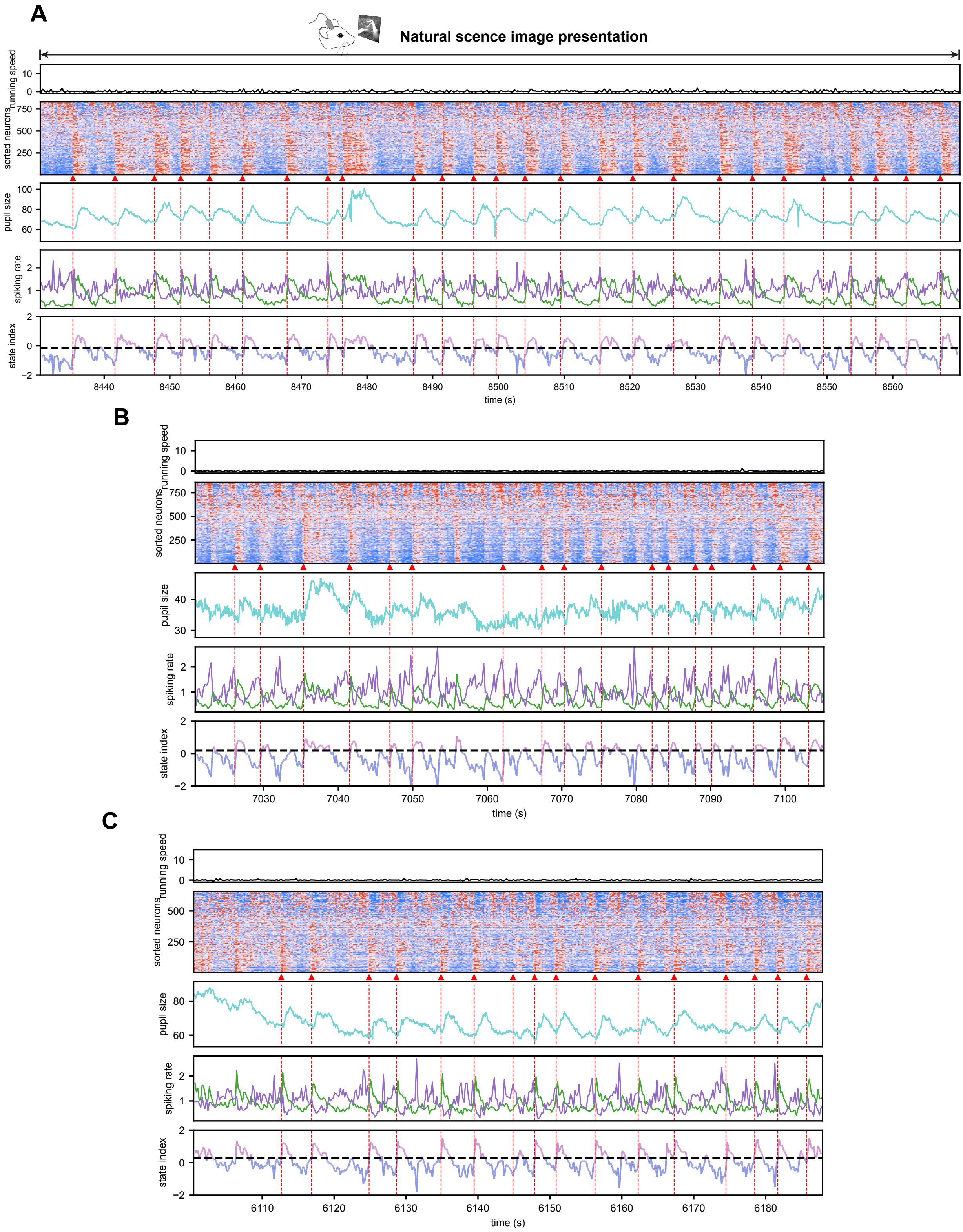

**Figure S3**. (A)–(C) Examples of spiking cascade during continuous natural scene image stimulation in the absence of running.

#### 2.4 Figure S4

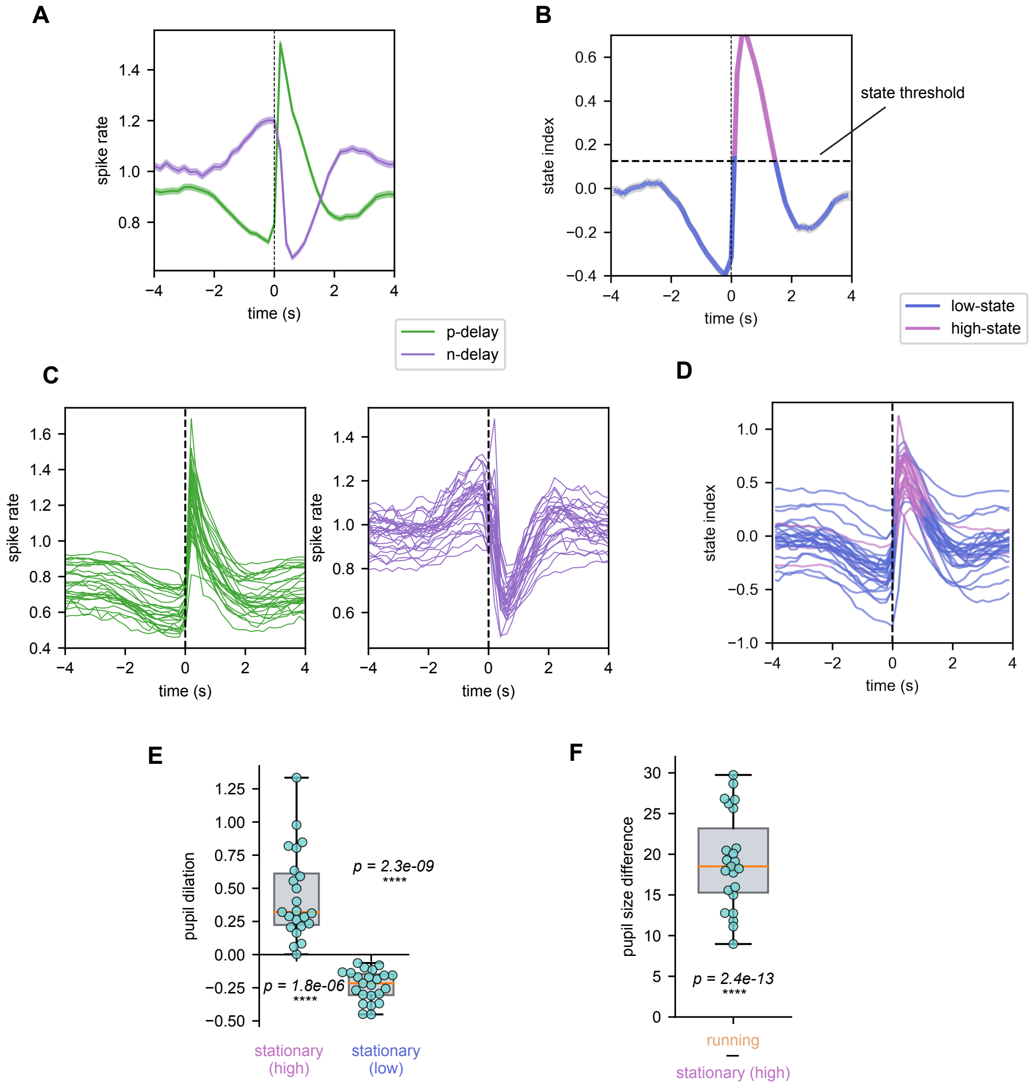

**Figure S4**. Spiking cascade is accompanied by seconds-scale arousal modulations. (A) Average activity of positive-delay neurons (green) and negative-delay neurons (purple) throughout the cascade cycle, shown for the representative mouse. (B) Averaged state index across the cascade cycle, illustrated for the representative mouse. The threshold distinguishing high and low states is indicated by a dashed line. (C) Average activity of positive-delay neurons (left, green) and negative-delay neurons (right, purple) throughout the cascade cycle across all mice. Each line represents a different mouse. (D) Averaged state index across the cascade cycle across all mice. Each line corresponds to a different mouse, and color denotes stationary high/low states. (E) Box plot presenting pupil dilation for both stationary-high and stationary-low states. Each dot represents an individual mouse. (F) Box plot demonstrating the difference in pupil size between running state and stationary-high state, with each dot representing a distinct mouse.

#### 2.5 Figure S5

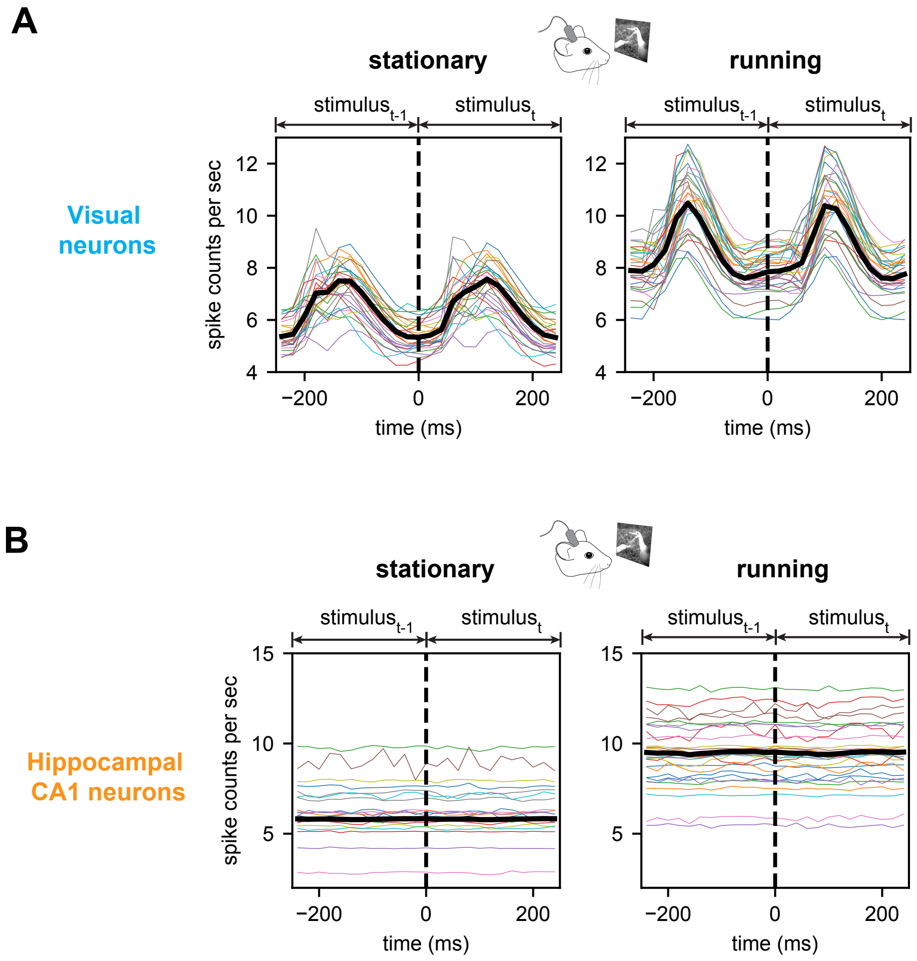

**Figure S5**. Neuronal responses to continuous image stimuli. (A) Mean activity of visual neurons displaying tuning responses to continuous image stimuli during both stationary and running periods. Each curve corresponds to a different mouse. (B) In comparison, the mean activity of hippocampal neurons remained independent of the continuous image stimuli.

#### 2.6 Figure S6

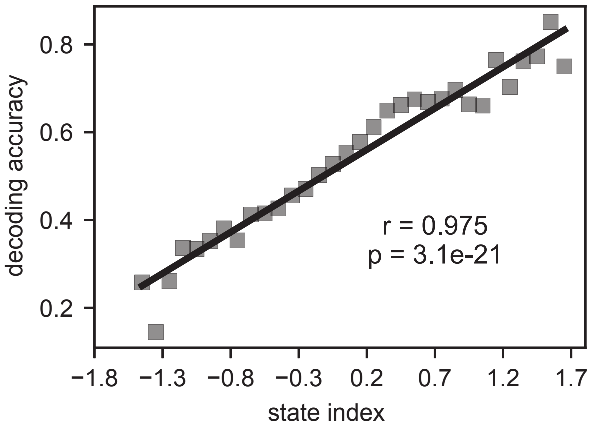

**Figure S6**. Linear relationship between state index and decoding accuracy averaged across all the mice during stationary periods. The linear relationship is measured by Pearson's correlation.

#### 2.7 Figure S7

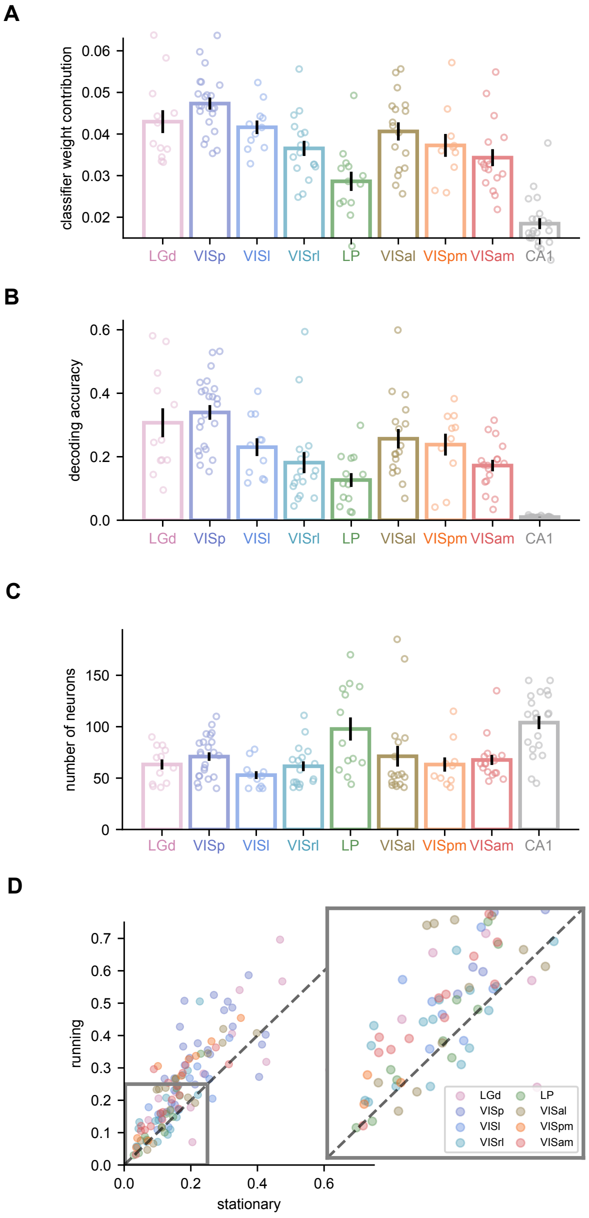

**Figure S7**. State dependent visual coding is evident in all visual areas. (A) Bar plot showing the averaged classifier weights importance of each brain region. Each dot represents a specific brain region in the corresponding mouse. (B) Bar plot illustrating the decoding accuracy of each brain region using data exclusively from that region. Each dot represents a region in the corresponding mouse. (C) Bar plot displaying the number of neurons in each region. Each dot represents a region in the corresponding mouse. (D) Changes in region-wise decoding accuracy between stationary and running conditions. Each colored dot represents a specific visual region from a single mouse.

#### 2.8 Figure S8

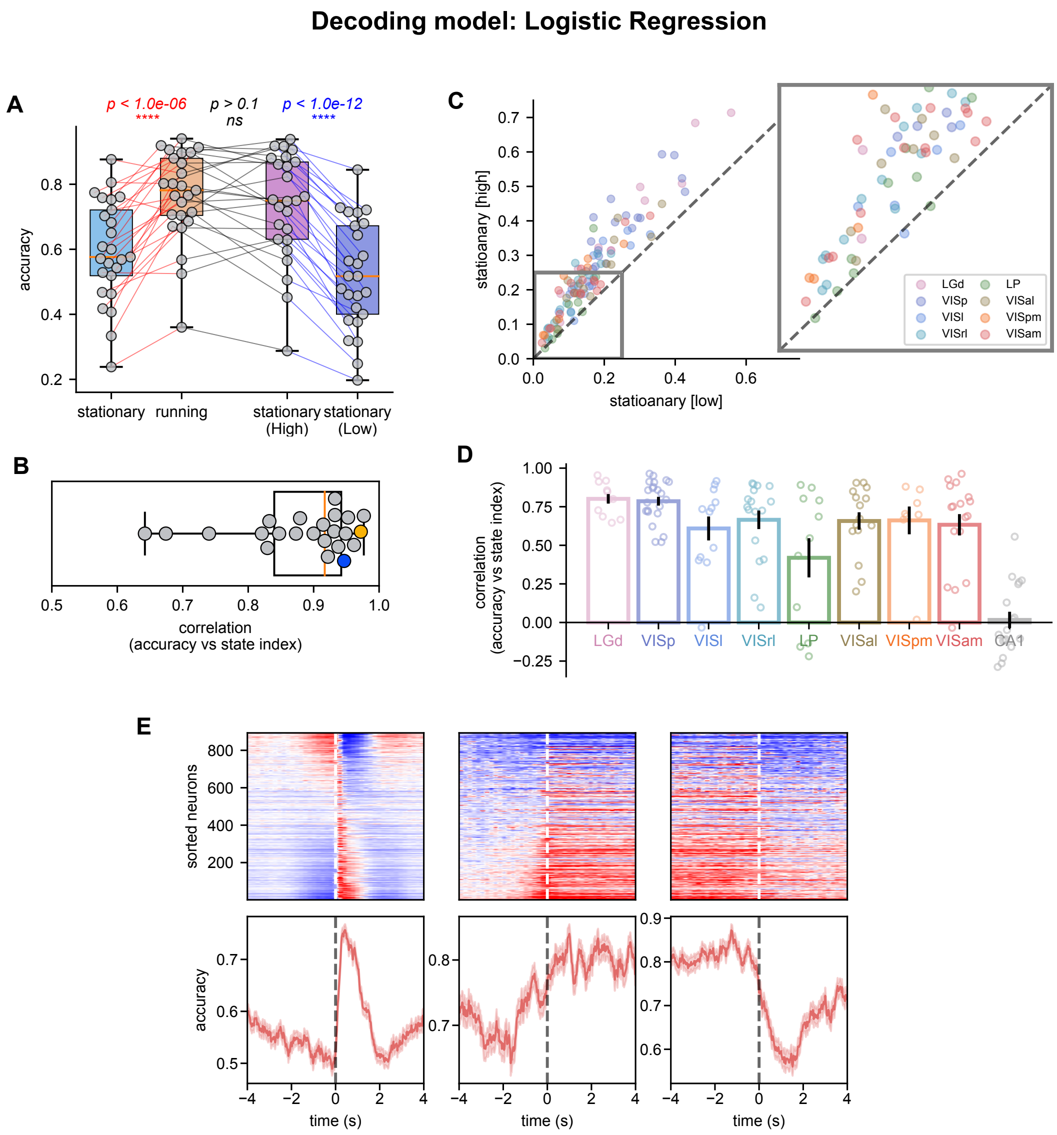

**Figure S8**. Decoding neural activities with logistic regression. (A) Box plot showing the decoding accuracy within-subject changes under different conditions. Each dot represents a mouse and pairwise t-test is used for significance test. (B) Linear relationship between state index and decoding accuracy is summarized in box plot for all mice where the yellow and blue dots represent the example mice correspondingly. The linear relationship is measured by Pearson's correlation. (C) Change in region-wise decoding accuracy between stationary-high and stationary-low state. Each colored dot represents visual region indicated by the color from a mouse. (D) Box plot showing the linear relationship between the state index and decoding accuracy for each brain region, similar to (C). Each dot represents a mouse with the corresponding region recorded. (E) Decoding accuracy across the 8-s cascade cycle (Left), running onset (middle) and offset (Right) from all 32 mice. Note the averaged cascade pattern is from the representative mice.

#### 2.9 Figure S9

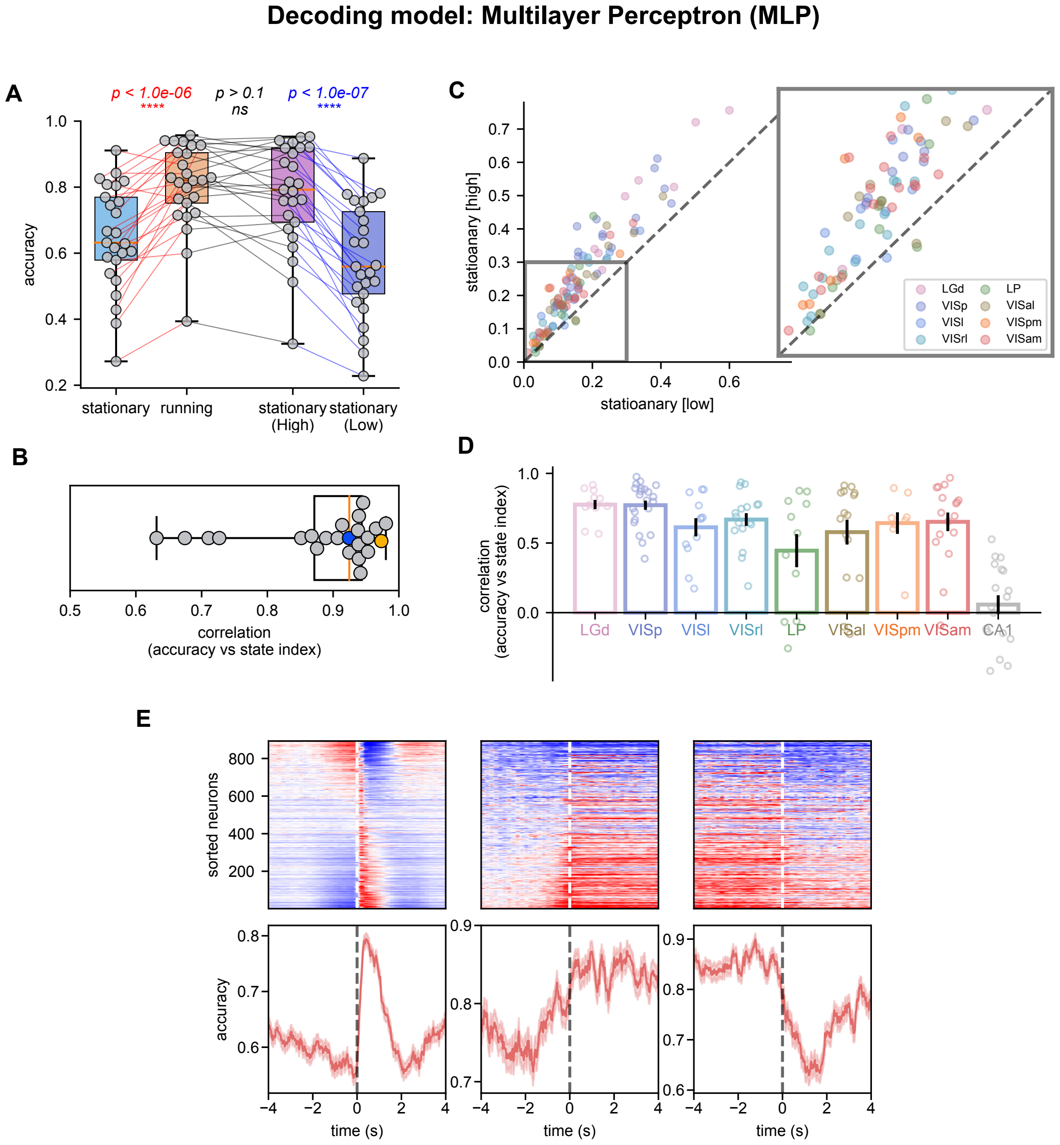

**Figure S9**. Decoding neural activities with multilayer perceptron (MLP). (A) Box plot showing the decoding accuracy within-subject changes under different conditions. Each dot represents a mouse and pairwise t-test is used for significance test. (B) Linear relationship between state index and decoding accuracy is summarized in box plot for all mice where the yellow and blue dots represent the example mice correspondingly. The linear relationship is measured by Pearson's correlation. (C) Change in region-wise decoding accuracy between stationary-high and stationary-low state. Each colored dot represents visual region indicated by the color from a mouse. (D) Box plot showing the linear relationship between the state index and decoding accuracy for each brain region, similar to (C). Each dot represents a mouse with the corresponding region recorded. (E) Decoding accuracy across the 8-s cascade cycle (Left), running onset (middle) and offset (Right) from all 32 mice. Note the averaged cascade pattern is from the representative mice.

#### 2.10 Figure S10

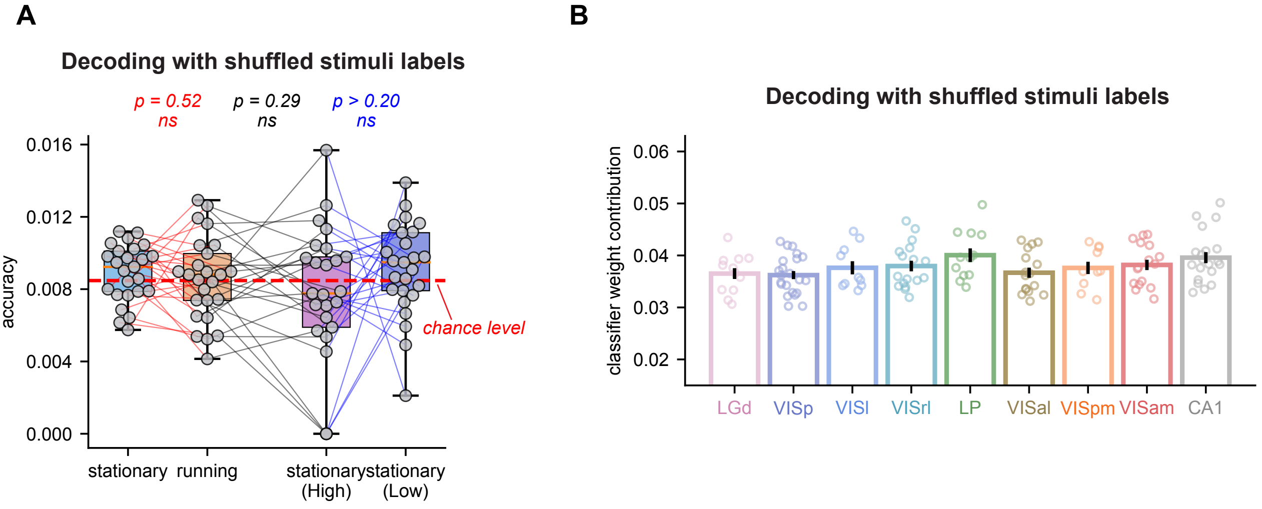

**Figure S10**. Decoding neural activities with shuffled stimuli labels. (A) Box plot showing the decoding accuracy within-subject changes under different conditions trained with shuffled stimuli labels. Each dot represents a mouse and pairwise t-test is used for significance test. (B) Bar plot showing the averaged classifier weights importance of each brain region with the classifier trained with shuffled stimuli labels. Each dot represents a region in the corresponding mouse.

#### 2.11 Figure S11

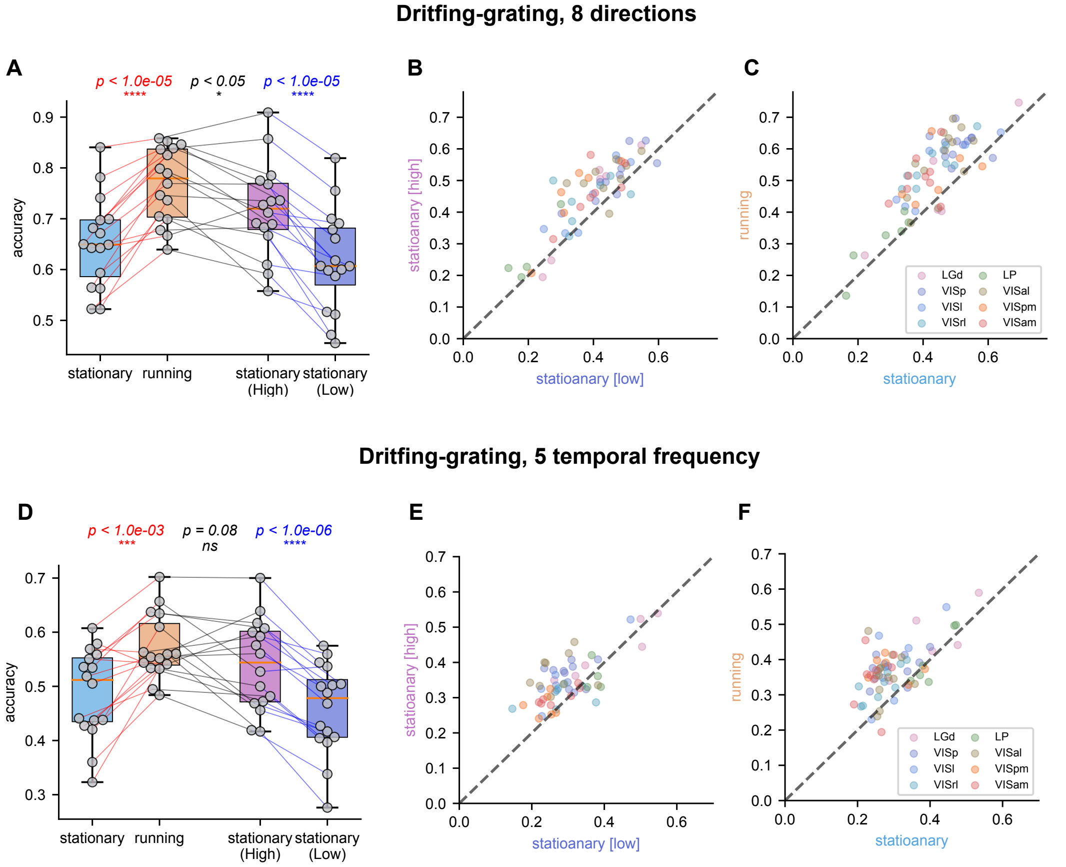

**Figure S11**. State-dependent population encoding of visual drifting-grating stimuli. (A) Box plot showing the accuracy of decoding drifting-grating direction within-subject changes under different conditions. Change in region-wise decoding accuracy of drifting-grating direction between stationary-high and stationary-low state (B) and between running and stationary (C). (D) Box plot showing the accuracy of decoding drifting-grating temporal frequency within-subject changes under different conditions. Change in region-wise decoding accuracy of drifting-grating temporal frequency between stationary-high and stationary-low state (E) and between running and stationary (F).

#### 2.12 Figure S12

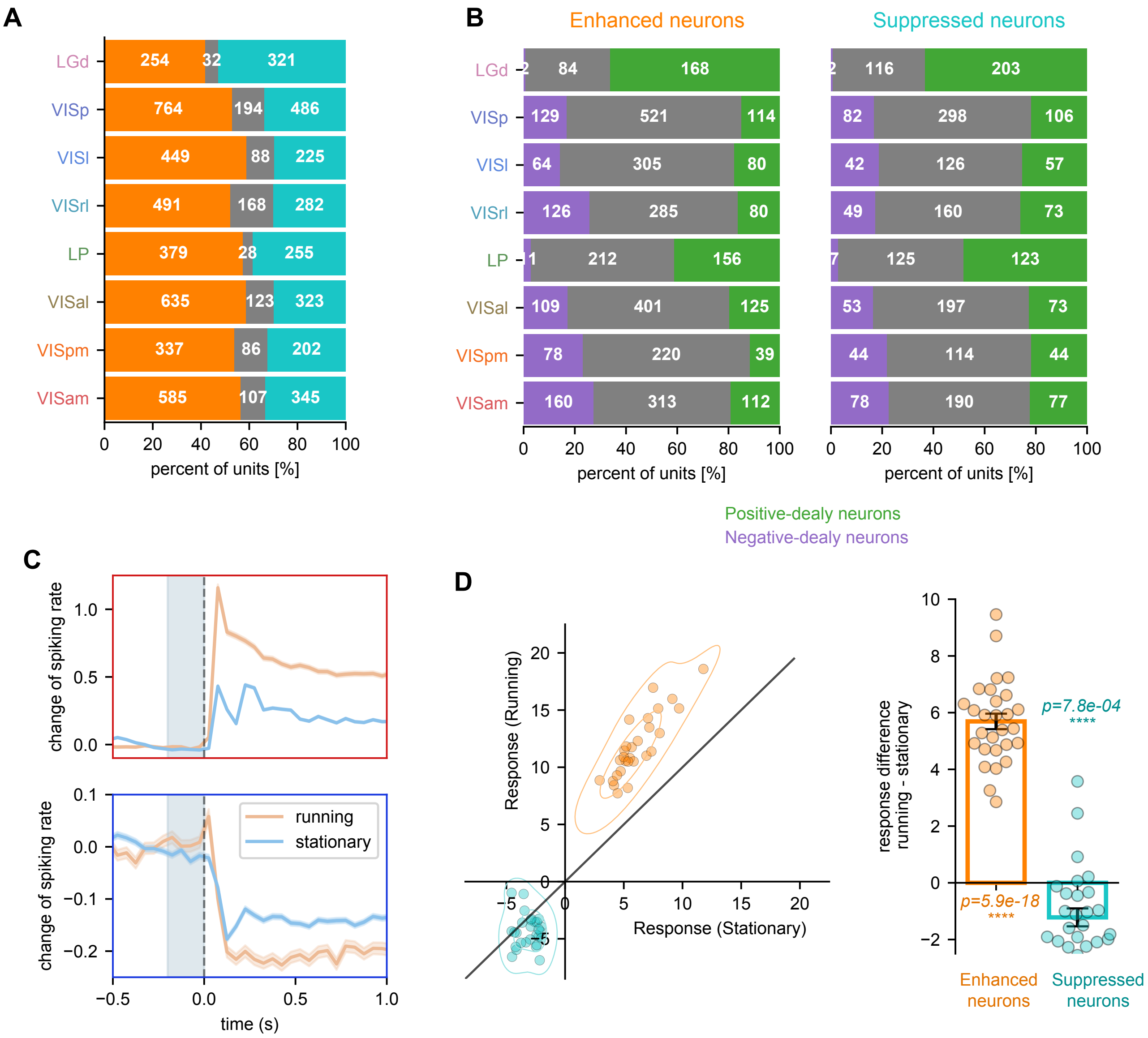

**Figure S12**. Single-neuron responses to Drifting-Grating stimuli. (A) Bar plot showing the percentage of drifting-grating responding neurons across visual regions, with neurons demonstrating exclusive positive responses represented in orange, exclusive negative responses in teal, and neurons with non-exclusive responses in gray.(B) Bar plot displaying the count of positive-delay neurons and negative-delay neurons within the population of enhanced neurons (left) and suppressed neurons (right). (C) The effects of pre-stimulus state (running / stationary) on the averaged stimulus-evoked response of enhanced neurons (top) and suppressed neurons (bottom) in the representative mouse. (D) Change in drifting-grating response between running and stationary. Single neuron response is quantified as the onset spiking rate (0 to 400ms) subtracted by the baseline (-800 to 0ms). Data is separated into two groups with orange color marked response of enhanced neurons and teal color marked that of suppressed neurons. Each dot within each group represents a mouse.

### 3 Materials and Methods

#### 3.1 Neuropixels Data

The present study conducted data analysis utilizing the Brain Observatory Neuropixel dataset obtained from the Allen Institute [1, 2]. This dataset comprises high-density extracellular neuron recordings of mice using Neuropixel probes. Each mouse was implanted up to six Neuropixel probes, which targeted the primary visual cortex (VISp) and five high-order visual cortical areas, namely, latero-medial area (VISl), anterol-ateral area (VISal), rostro-lateral area (VISrl), postero-medial area (VISpm), and antero-medial area (VISam). The silicon probes were inserted to a depth of up to 3.5mm into the brain, enabling the recording of spiking activity within two visual thalamic nuclei, i.e., the lateral posterior nucleus (LP) and the lateral geniculate nucleus (LGN), as well as other regions that the probes traversed, such as the hippocampus (Fig. 1A and 3A).

We focused on the experiments from all 32 mice performed with "Brain Observatory 1.1" stimulus set containing various types of visual stimulus as shown in Fig. S1A. We specifically narrowed our analysis to two types of stimuli: natural images and drifting gratings. During each experiment, the natural image and drifting grating stimuli were presented separately in three distinct sessions.

Regarding the natural image stimuli, a continuous sequence of images was presented in each session, with each image maintained for a duration of 250ms. Across the entirety of the experiment, a total of 118 distinct image stimuli were employed, each being repeated 50 times and randomly delivered throughout the three sessions (as illustrated in Figure 1B).

For the drifting gratings, stimuli were displayed with a spatial frequency of 0.04 cycles/deg, 80% contrast, 8 directions (0°, 30°, 60°, 90°, 120°, 150°) and 5 temporal frequencies (1, 2, 4, 8 and 15 Hz). Across all three drifting grating sessions, each condition was repeated 15 times.

#### 3.2 Mice Exclusion

We did not confine the analysis to a small subset of mice; rather, distinct subsets were employed for various analyses to fully exploit the dataset. Initially, three mice were excluded due to insufficient stationary periods during task sessions (less than 10% of the time), resulting in a remaining cohort of 29 mice.

- For the pupil size analysis, six mice lacking pupil data were excluded, yielding a cohort of 23 mice for the analysis.
- In the natural scene visual stimuli analysis, the objective was to compare decoding accuracy across three conditions: stationary-high, stationary-low, and running periods. Given a total of 5900 samples for 118 images with 50 repeats, mice with an insufficient number of samples ($n \leq 150$) for each condition were excluded, leading to the removal of nine mice and leaving 20 mice for analysis.
- In the drifting-gratings analysis, aimed at comparing the response of positive-responding and negative-responding neurons across three conditions, one mouse lacking stationary periods during the drifting-grating session was excluded. This resulted in a final cohort of 28 mice included in the analysis.

Removal details are shown in Table 1.

#### 3.3 Spiking Cascade Detection

Processed neural spike data from the Allen Neuropixel dataset were utilized in our analysis. The neural spike data were sorted with Kilosort2 pipeline [3]. Further information regarding the processing of the spike data can be found in the white paper of the dataset.

Following prior studies [4, 5] to identify spiking cascade events, we first divided time evenly into 200ms time bins and calculated spike rates by counting the number of spikes within each bin. Then we applied a delay-profile decomposition method to extract the relative temporal relationship among a group of neurons in a data-driven manner. Briefly, we identified candidate neural events by segmenting the spiking rate data based on the troughs of the filtered global mean spiking rate (low-pass, 0.25Hz). For each neural event $k$, we derive a delay profile vector $\boldsymbol{d}_k$ representing the temporal phase of each neuron within that event.

$$\boldsymbol{d}_k = \begin{pmatrix} t_{1k} & t_{2k} & \cdots & t_{Nk} \end{pmatrix}^T \tag{1}$$

where $t_{i,k}$ representing the temporal centroid of the firing rate within the time segment $k$ and $N$ represent the total number of neurons in the recording. A delay profile matrix $\boldsymbol{D}$ representing all of the potential temporal relationships among the neurons can thus be constructed by combining the delay profiles for all $M$ time segments.

$$\boldsymbol{D} = \begin{pmatrix} \boldsymbol{d}_1 & \boldsymbol{d}_2 & \cdots & \boldsymbol{d}_M \end{pmatrix} \tag{2}$$

$$= \begin{pmatrix} t_{11} & t_{12} & \cdots & t_{1N} \\ t_{21} & t_{22} & \cdots & t_{2N} \\ \vdots & \vdots & & \vdots \\ t_{M1} & t_{M2} & \cdots & t_{MN} \end{pmatrix} \tag{3}$$

Singular value decomposition (SVD) was then applied to the delay matrix $\boldsymbol{D}$ with principle component $\boldsymbol{u}^*$ defined as the principal delay profile representing the major sequential organization among the group of recorded neurons. We then use the principal delay profile as a template to match the delay profile of candidate events. A candidate event that has a significant similarity score ($p < 0.001$) measured with Pearson's correlation was considered as a cascade event in our study.

#### 3.4 State Index

##### 3.4.1 Positive-delay and Negative-delay Neurons

In the context of a cascade cycle, it is evident that the behavior of neurons is contingent upon their specific activation phase. In light of this observation, we considered two discrete categories of neurons based on their activation phase within the spiking cascade cycle. These categories are denoted as the negative-delay neurons (depicted by purple symbols in Figure 1C) and the positive-delay neurons (represented by green symbols in Figure 1C).

The negative-delay neurons exhibited a gradual recruitment during the initial phase of the cascade, followed by a rapid transition to the subsequent activation of positive-delay neurons. The latter were subsequently discharged in a sequential manner during the later phase of the cascade. Formally, the negative-delay neurons $\mathcal{N}$ and positive-delay neurons $\mathcal{P}$ were defined as those whose mean delay value across all the candidate events, was significantly ($p < 0.001$) lower or higher than zero respectively.

##### 3.4.2 State Index Definition

Given the prominent role of the two neuron populations in regulating brain states (Fig. 1C, S4), we derived a state index $s(t)$ by quantifying their relative activities for each time $t$:

$$s(t) = \gamma \overline{f_{\mathcal{P}}(t)} - (1-\gamma)\overline{f_{\mathcal{N}}(t)} \tag{4}$$

$$\text{with } \gamma = \frac{\sigma_{\mathcal{N}}}{\sigma_{\mathcal{N}} + \sigma_{\mathcal{P}}} \tag{5}$$

where $\overline{f_{\mathcal{P}}(t)}$, $\overline{f_{\mathcal{N}}(t)}$, $\sigma_{\mathcal{P}}$, $\sigma_{\mathcal{N}}$ represent mean firing rate of positive-delay neurons, negative-delay neurons, and standard deviation of the mean firing rate of positive-delay neurons and negative-delay neurons respectively.

The distribution of the state index exhibits a bimodal pattern, as shown in Figure 1E. Consequently, a two-class Gaussian mixture model was utilized to accurately model this bimodal distribution of the state index. The separation of these modes was achieved through the application of a threshold that corresponds to the optimal decision boundary. Specifically, periods characterized by state index values exceeding the threshold were categorized as stationary-high states, while periods featuring state index values below the threshold were designated as stationary-low states.

#### 3.5 Neural Population Decoding Analysis

To assess the sensory information encoded within the neuron population, we conducted an analysis of spiking data using nature scene image stimulation. In this protocol, a sequence of images was presented continuously, with each image stimulus having a duration of 250ms. For each image stimulus, we defined the neural code as a vector in which each entry corresponds to the count of spikes from a specific neuron, measured within a 200ms time window following the onset of the stimulus. For example, the neural code for the $i$th stimulus is

$$\boldsymbol{SR}_i = \begin{pmatrix} f_{1i} & f_{2i} & \cdots & f_{Ni}, \end{pmatrix} \tag{6}$$

where $f_{ki}$ denotes the $k$th neuron spike count within 200ms period following the $i$th stimulus onset.

We then developed a method to quantify the degree of visual information encoded in the population neural code for each stimulus. The image stimuli were shuffled and divided into five groups, ensuring equitable distribution of stimuli classes. Within each group, there were 10 repetitions for every class, and special attention was given to prevent any overlap of stimuli across these groups. For each iterative step, we designated one group as the test dataset, while the remaining four groups served as training data. In the training phase of each iteration, we employed a Support Vector Machine (SVM)-based decoder. This decoder was trained to establish a link between the population neural codes and the corresponding stimulus classes. Subsequently, we employed this trained decoder to make predictions on the test data. This process was repeated for all five iterations, and as a result, each stimulus was assessed and categorized as either correctly classified or misclassified. The decoding accuracy was computed as the average rate of accurate classification across all stimuli:

$$acc = \frac{1}{n}\sum_{k=1}^{n} C_k \tag{7}$$

$$\text{with } C_k = \begin{cases} 1 & \text{if k is correctly classified} \\ 0 & \text{if k is misclassified,} \end{cases} \tag{8}$$

where $n$ is the number of stimuli in consideration. In this manner, the decoding accuracy can be evaluated for different states or conditions, such as immobile and locomotion states.

In addition to natural scene image stimuli, we also conducted the same decoding analysis for drifting-grating stimuli to estimate stimulus direction and temporal frequency (Fig.S11). To validate the robustness of the analysis, we examined various decoder models, i.e., logistic regression (Fig. S8) and three-layer multilayer perceptron with 512 hidden units (Fig. S9). To validate the integrity of our methodology, we conducted a control analysis using the exact same approach. The only distinction was that we shuffled the stimulus identities (labels) randomly, as shown in Figure S10. This control analysis helped us ascertain the robustness of our results.

To evaluate the impact of the state index on decoding accuracy, we grouped the trials based on their respective state index values and subsequently calculated the mean decoding accuracy for each grouping. Within our study, we maintained group separations using a differential of 0.1 in terms of the state index values.

#### 3.6 Neuron Evoked Responses Analysis

We analyzed single neuron encoding of external stimuli with spiking data from drifting-grating session as it contains a 1 second baseline period before every stimulus, providing suitable noise control. Drifting-grating responding neurons were defined as those exhibiting a statistically significant difference in spiking rate during the stimulation period (0-600ms post stimulus onset) compared to the baseline period (-800-0ms), as determined by a paired t-test with a significance level of $p < 0.001$.

These drifting-grating responding neurons could either be significantly enhanced or suppressed by the presented stimuli. Accordingly, we classified neurons as "enhanced" or "suppressed" based on whether they displayed an increase or decrease in spiking rate, respectively, during stimulus presentation.

We next investigated how pre-stimulus brain state affects single neuron response. The neuron evoked response was quantified by the spiking rate difference between the stimulation onset (0-400ms) and baseline periods (-800-0ms). As single neuron activity might be confounded by the ongoing state-dependent dynamics, we focused on the drifting-grating responding neurons without those significantly modulated by the cascade, i.e., the positive-delay and negative-delay neurons (Fig. S12B). The neural responses elicited by each stimulus separately for both enhanced neurons and suppressed neurons are defined as:

$$e^i_{\mathcal{E}_i} = \frac{1}{|\mathcal{E}_i|} \sum_{k \in \mathcal{E}_i} f^i_k \tag{9}$$

$$e^i_{\mathcal{S}_i} = \frac{1}{|\mathcal{S}_i|} \sum_{k \in \mathcal{S}_i} f^i_k \tag{10}$$

where $\mathcal{E}_i$, $\mathcal{S}_i$ represent the set of enhanced neurons and suppressed neurons of the stimulus $i$, $e^i_{\mathcal{E}_i}$, $e^i_{\mathcal{S}_i}$ represent the evoked response for stimulus $i$ from the two neuron sets, and $|\cdot|$ denotes the size of the set. The pre-stimulus state was defined as the state index during the 200ms pre-stimulus period.

#### 3.7 Local Field Potential Processing

Delta power and gamma power were computed for local field potentials (LFPs) across all recorded channels. To calculate delta power, a band-pass filter (1-4 Hz) was applied to the LFP signal of each channel, followed by rectification and lowpass filtering ($< 0.72$ Hz, corresponding to $\pi$ cycles of the mean band-pass frequencies). Similarly, gamma power was extracted using a comparable procedure, with the exception of applying a band-pass filter (55-65 Hz) and setting the low-pass filter cutoff frequency to 19 Hz with respect to $\pi$ cycles.

Hippocampal sharp wave ripples (SWRs) are brief, high-frequency oscillations (110-200Hz) that can be observed in the local field potential (LFP) recorded from hippocampal recording sites. For ripple detection in this study, we employed an offline method [4, 6] utilizing the LFP signal (1250 Hz) captured from the hippocampal CA1 region. The identification of ripple events was conducted individually for each CA1 recording site (channel), resulting in robust and extensively overlapping ripple detection across the channels. To consolidate the detection outcomes from various channels, a criterion was imposed: a detected ripple event was deemed valid only if it was identified in more than 40% of the CA1 channels.

#### 3.8 Behavior Data

##### 3.8.1 Stationary and Running State

Throughout the recording sessions, the mice exhibited significant amounts of both running and stationary periods. To delineate these periods, we first applied a low-pass filter (a 3rd-order Butterworth filter with a cutoff frequency set at 2Hz) to the running speed data extracted from the dataset. This filtration aimed to eliminate high-frequency noise. Negative running speeds were attributed solely to sensor noise [4]. Consequently, a threshold was determined at the 0.05 percentile of the absolute values of the negatively filtered speed values. Any time point at which the filtered running speed exceeded this threshold was categorized as a non-stationary point.

Given that short stationary periods were interspersed with non-stationary periods, our selection process focused exclusively on stationary periods that persisted for a duration exceeding 20 seconds. Additionally, these stationary periods were required to have a minimum gap of 3 seconds between identified stationary and non-stationary periods to mitigate potential boundary effects. Similarly, running periods were identified using the same approach, but the selection was confined to non-stationary periods lasting longer than 0.5 seconds and a minimum time gap of 3 seconds was mandated between stationary and running periods.

##### 3.8.2 Pupil Size

The eye-tracking data from the dataset were pre-processed with a set of parameters pre-computed at sampling rate of 30 Hz. We used pupil diameter as the arousal index in our analysis and it is defined as the mean of the pupil height and width. To eliminate high-frequency noise, the pupil diameters were subjected to low-pass filtering with a cutoff frequency set at 2 Hz.
